# Longitudinal computer vision-based behavioural phenotyping of group-housed dogs for translational and welfare research

**DOI:** 10.64898/2026.11.19.767794

**Authors:** Miho Sato, Preami Uthayavathanan, Juval Gutknecht, Andrea Campa, Thomas Lorimer, Paula Vazquez-Pianzola, Kazhal Almasi, Lorena Item, Dominik N’Gouan, Oliver Sturman, Henning Richter

**Affiliations:** Diagnostic Imaging Research Unit, Clinic for Diagnostic Imaging, Department of Clinical Diagnostics and Services, Vetsuisse Faculty, University of Zurich, Switzerland; VSF Stiegenhof, Vetsuisse Faculty, University of Zurich, Switzerland; 3R Hub, ETH Zurich, Switzerland; IT support, Vetsuisse Faculty, University of Zurich, Switzerland

**Keywords:** Dogs, Behavioural analysis, re-identification, individual behavioural profiling, Group-housing, Posture recognition, Phenotyping, Computer vision, YOLO, Deep learning, Multi-model integration, Circadian rhythm, Animal welfare

## Abstract

Continuous behavioural monitoring is essential for assessing animal welfare and individual behavioural variation, yet traditional observation-based methods are labour-intensive and prone to bias. Here, we developed a modular framework with computer vision for automated behavioural profiling of group-housed Beagles, integrating You Only Look Once (YOLO)-based models for pose estimation, individual identification, and enrichment detection. A ceiling-mounted infrared camera system enabled uninterrupted monitoring under standard kennel conditions. The identity-independent YOLO-POSE model achieved 99% precision, 97% recall, and an mAP50 of 0.98, while the YOLO classification model reached 95% accuracy for individual identification without external markers. Combining posture, identity, and environmental information enabled automated quantification of spatial use, locomotor activity, social interactions, corridor-oriented behaviour, drinking behaviour, and enrichment use. Overall, this framework captured distinct behavioural profiles of individual dogs from the same litter housed in the same kennel, which can be used to assess animal welfare and to evaluate environmental enrichments objectively. Together, we demonstrate a scalable and non-invasive approach for continuous individual behavioural profiling of group-housed dogs for computational ethology and translational research. By separating pose estimation from identity assignment, the framework enables flexible deployment across changing social groups and facilitates application to previously unseen individuals with minimal training.

## Introduction

Animal behaviour is a sensitive indicator of how animals perceive and respond to their physical and social environment, as well as their physical and mental state (Broom, 1991; M. S. Dawkins, 2003; Mellor, 2016). Therefore, behaviour serves as a central outcome measure across behavioural, welfare, and translational research (Duncan & Dawkins, 1983; Broom, 1991; Krakauer et al., 2017). Animal behaviour can exhibit individual differences that are repeatable and consistent over time and across situations, and that may be biologically meaningful (Réale et al., 2007; Bell et al., 2009; Stamps, 2016). Traditional behavioural assessments include direct observation, questionnaires, and ethograms (van den Berg et al., 2003; Van Der Waaij et al., 2008; Thallmair et al., 2026). While these methods remain valuable to assess animal welfare (M. Dawkins, 2004; Bateson & Martin, 2021), they are inherently limited by their reliance on human observation, making them labour-intensive, temporally limited, and susceptible to inter-observer variability (Altmann, 1974; De Passillé & Rushen, 2005; Bateson & Martin, 2021). Moreover, the presence of observers can influence animal behaviour (Crofoot et al., 2010; Sherwen & Hemsworth, 2019; Allan et al., 2020). In welfare science, this is particularly consequential: gradual behavioural changes associated with chronic stress, pain, or compromised affective states may be imperceptible within short observation windows, meaning that welfare-relevant changes are among the most likely to go undetected by conventional methods (Barnard et al., 2016; Veissier et al., 2019). Although sensors provide another approach to behavioural assessment (Whitford & Klimley, 2019), their deployment is constrained by predefined measurement parameters, the potential to induce stress responses, and limited battery life (Gerencsér et al., 2013; Stygar et al., 2021; Chelotti et al., 2024). As a result, these approaches have difficulty capturing gradual, continuous, and long-term behavioural dynamics.

Computer vision (CV) and machine learning have transformed behavioural research. First, by automating object detection, tracking, and behavioural classification from images and videos, CV enables high-throughput behavioural analysis (Anderson & Perona, 2014; Egnor & Branson, 2016; Fazzari et al., 2025). Second, by decoupling data collection from scoring, CV reduces human bias and alleviates constraints imposed by predefined measurement schemes, allowing more flexible post hoc analysis. Third, the non-invasive nature of CV enables continuous and repeated recording of behaviour, which is crucial for investigating behavioural dynamics across different timescales and individual differences. Finally, by providing fine-grained spatiotemporal data for each individual, CV facilitates the detection of previously uncharacterised behavioural patterns that may be overlooked using conventional observational approaches. Among different feature-extraction paradigms, keypoint-based pose estimation has emerged as an informative framework for representing animal posture (M. W. Mathis & Mathis, 2020), as it encodes body configurations in contrast to object detection using bounding boxes (BBOX) and instance segmentation using masks. Beyond simple measures of activity and location, pose-estimation methods offer the potential to quantify posture, movement, and behavioural states, thereby expanding the scope of automated behavioural analysis. The ability to quantify posture and movement may facilitate the identification of behavioural patterns that cannot be inferred from spatial location alone. For example, DeepLabCut (DLC) enables markerless pose estimation through transfer learning-based keypoint detection in single- and multi-animal settings (A. Mathis et al., 2018; Lauer et al., 2022). DLC can be extended to support multi-animal tracking with individual re-identification (Re-ID). Similarly, Social LEAP Estimates Animal Poses (SLEAP) provides a fully integrated framework for multi-animal pose estimation with built-in identity assignment and tracking of identities across frames in social environments (Pereira et al., 2022). Re-ID may be particularly valuable in species that exhibit substantial individual behavioural variation (Réale et al., 2007; Bell et al., 2009). However, robust long-term behavioural monitoring in complex group-housing environments remains challenging, particularly because of ambiguities in identity tracking under dynamic group composition. While pose estimation remains applicable without re-annotation when group composition changes, identity consistency over time can become ambiguous, limiting the long-term use of trained models.

Dogs occupy a unique position as both a welfare-relevant species in their own right and a valuable translational model (Hubrecht et al., 1992; Rooney et al., 2009). As companion and managed animals, dogs share physical and social environments with humans, while their naturally occurring diseases often parallel human conditions, supporting their value as translational models (Hytönen & Lohi, 2016; Ostrander et al., 2019). Compared with traditional rodent models, their physiological, cognitive, and behavioural similarities to humans have further contributed to their increasing use in studies of neurological, oncological, cardiovascular, and other complex disorders (Lindblad-Toh et al., 2005; Hasiwa, 2011). In addition to their value as translational models, dogs housed in kennels or research facilities represent an important population for behavioural and welfare research. In particular, group-housed individuals share a common environment yet differ consistently in their behavioural responses to housing conditions, social partners, and husbandry routines (Sih et al., 2004; Réale et al., 2007; Bell et al., 2009; Dingemanse et al., 2010), and this variation cannot be adequately captured by population-level summaries (Hertel et al., 2020). Such individual differences in spatial preferences, activity patterns, and responses to environmental change may reflect meaningful variation in how individual animals experience and adapt to their managed environment (Gartner, 2015; Langbein & Nawroth, 2022). Quantifying these differences requires behavioural data collected continuously and at the level of the individual, across the full range of daily conditions, including rest, activity, and responses to routine husbandry events.

In canine research, automated behaviour tracking, pose estimation, and behaviour classification approaches have been applied to quantify movement (Zamansky et al., 2019), posture (Barnard et al., 2016; Ferres et al., 2022; Yu & Choi, 2022; Shooter et al., 2024), behavioural states (Barnard et al., 2016; Chavez-Guerrero et al., 2022; Eberhardt et al., 2026), and facial expressions (Waller et al., 2013). However, existing approaches have not yet enabled continuous, scalable, and individual-level behavioural monitoring of dogs in complex group-housing environments.

To address these limitations, we developed an objective framework for continuous individual behavioural profiling of group-housed dogs using real-time You Only Look Once (YOLO) models (Redmon et al., 2016; Ultralytics, 2024; Jocher et al., 2026). A YOLO-POSE model was adapted to estimate the posture of individual dogs, a trained YOLO classification model (YOLO-CLS) was used to determine individual identity, and a YOLO oriented BBOX (OBB) model (YOLO-OBB) was trained to detect enrichment features, including beds, gateways, and water bowls. Integrating these models enabled the quantification and visualisation of spatial occupancy, locomotor activity, inter-individual relationships, responses to environmental modifications, and interactions with specific resources. Sequential behavioural patterns further revealed distinct time-of-day-dependent behavioural states without predefined behavioural labels. The proposed framework provides a scalable approach for continuous behavioural phenotyping of group-housed dogs, enabling objective comparison of individuals over time and under different housing conditions, enrichment strategies, and husbandry conditions. By quantifying behavioural variation and responses to environmental change, this approach may support evidence-based evaluation of welfare-related management practices while facilitating long-term behavioural monitoring in kennel environments.

## Results

### System setup for continuous behavioural monitoring

To establish continuous behavioural monitoring of group-housed Beagles in a translational research kennel, a camera system was installed centrally on the ceiling of each kennel (Fig. 1a). Infrared illumination enabled video acquisition during dark periods while maintaining standard housing conditions. To minimise visual occlusions and improve animal tracking, the kennel interior was modified by replacing bench-mounted beds with elevated beds while remaining fully compliant with relevant animal housing regulations. This configuration enabled uninterrupted video recording under varying environmental lighting conditions, including daylight fluctuations and nocturnal periods (Fig. 1b and c).

**Fig. 1.**
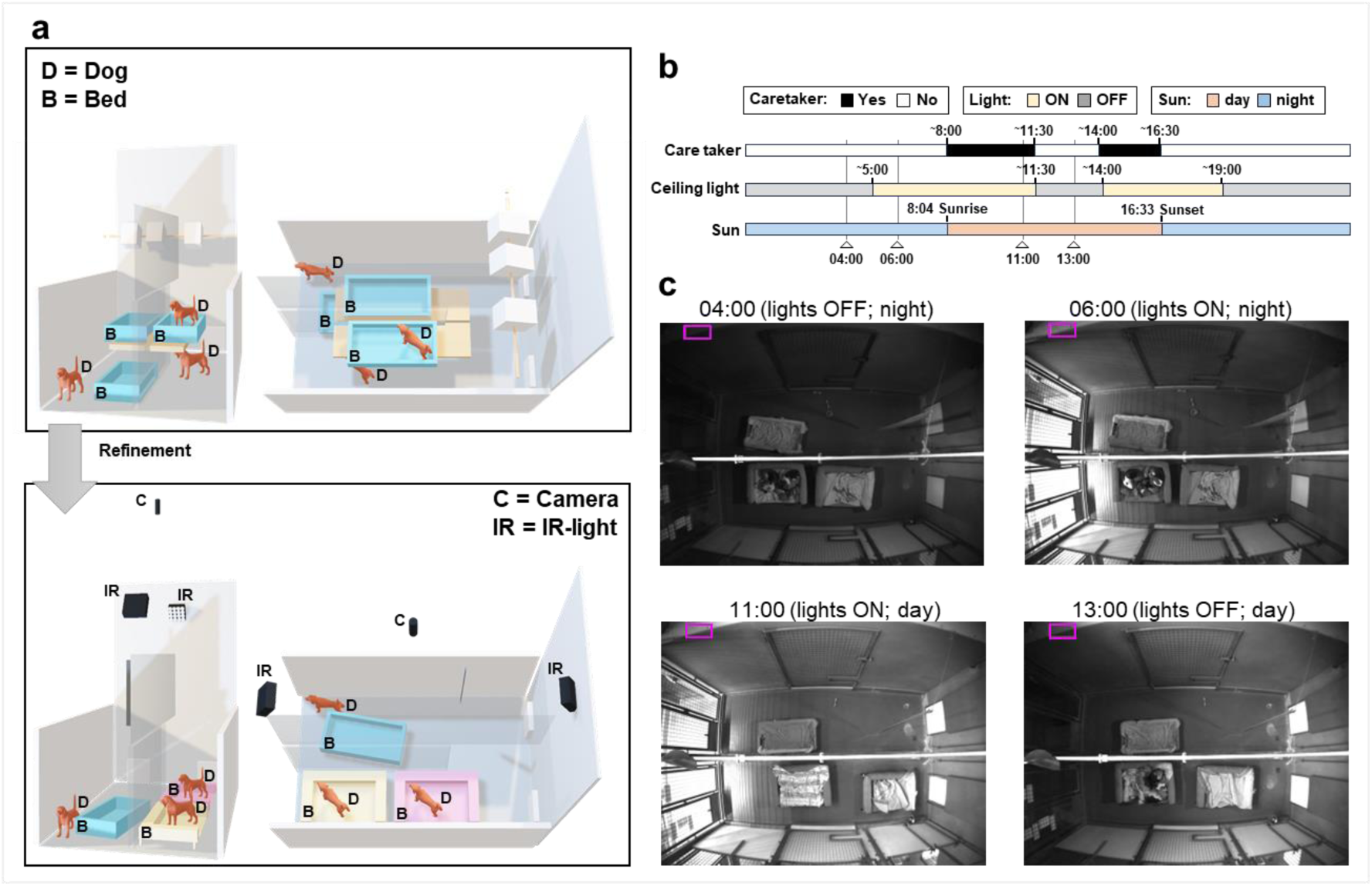
Transformation of housing kennels installing cameras. **a)** Three-dimensional representation of an indoor kennel showing key elements used for data acquisition. The original (a) and the final (b) furniture arrangements are shown from the side (left) and top (right) views. Original beds are shown in blue and new elevated beds in yellow. **b)** Daily schedule of dogs on a representative day. **c)** Representative video frames across different times of day indicated with triangles in b. The magenta rectangle indicates the region of interest (ROI) used to analyse brightness in the kennel.

### Quality control of video recordings

Beyond animal movement, video recordings inherently capture environmental variation, which can influence animal behaviour. Therefore, non-ML CV analysis was conducted to extract such variation (Supplementary Fig. 1). Mean absolute error (MAE) analysis of consecutive frames showed pronounced variation during daytime periods and lower values during phases when ceiling lights were turned off and caretaker activity was absent (Fig. 2a). Brightness measurements confirmed the scheduled lighting conditions, showing abrupt increases and decreases corresponding to light-on and light-off events, respectively. Sunrise and sunset during the light-on period were colocalised as gradual changes in brightness (Fig. 2b). Analysis of door status identified periods when kennel doors were open (Supplementary Fig. 2 and Fig. 2c).

**Fig. 2.**
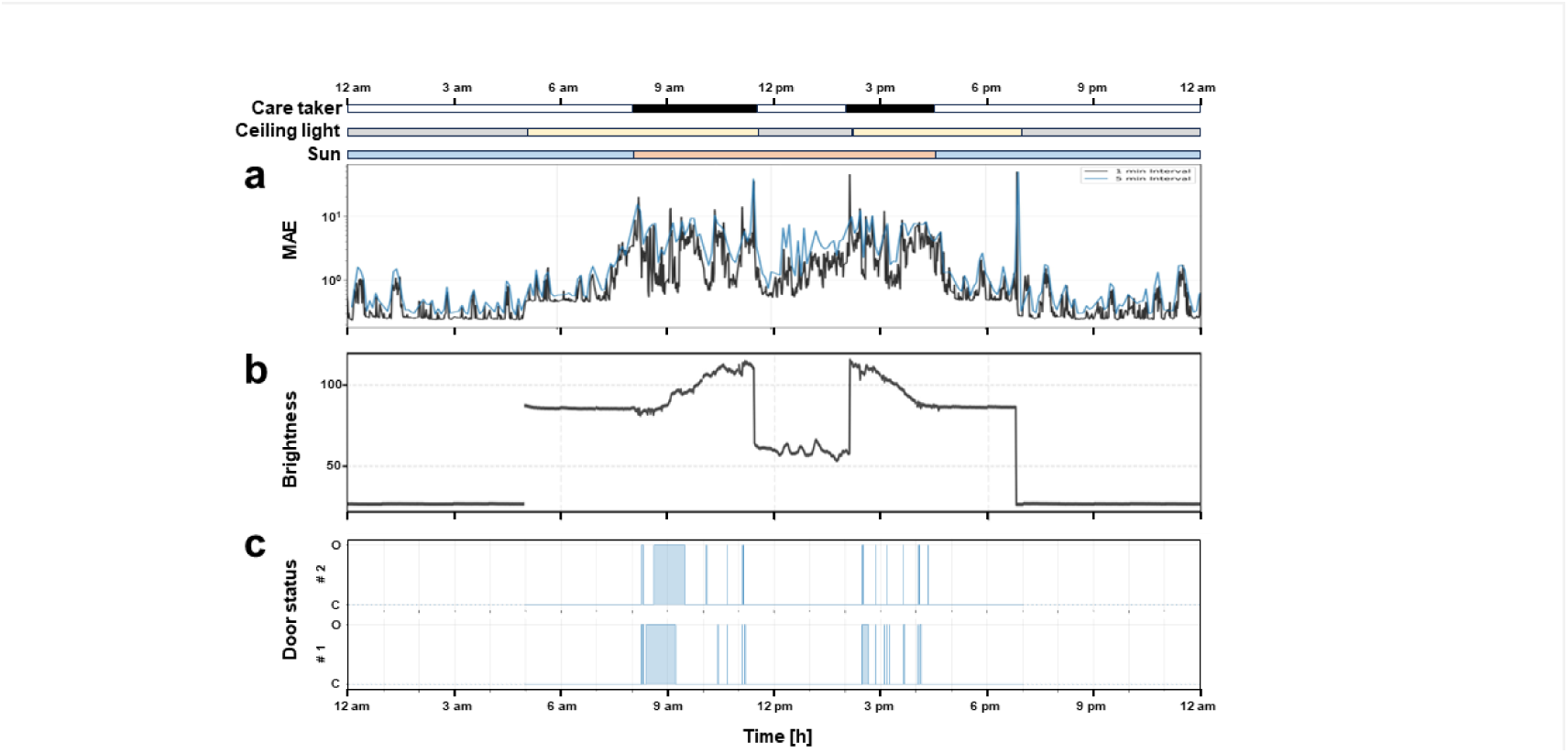
Non-ML CV analysis for environmental variation. **a)** MAE as a proxy for pixel changes computed per minute (black) or per 5 minutes (blue). **b)** A proxy for brightness detected as average pixel intensity (magenta rectangle) within the top-left region of the kennel wall as. The pixel area used for the analysis is shown in Fig.1c. **c)** Status of kennel doors on the corridor: open (O) or closed (C). The blue lines show the status of the doors. The blue boxes indicate closed status of each door.

### Pipeline design

Although individual Beagles displayed distinguishable coat patterns (Fig. 3a and b), developing an identity-specific pose estimation model would substantially reduce scalability and flexibility. To overcome this limitation, a modular dual-model strategy was developed consisting of: (i) a trained YOLO-POSE model for identity-independent body part detection and (ii) a trained YOLO-CLS model for individual classification (Fig. 3c and Supplementary Fig. 1). Female dogs were annotated with 29 digital keypoints to train the YOLO-POSE model (Fig. 3d, Supplementary Fig. 3, and Supplementary Fig. 5a and b), and YOLO-POSE model achieved the lowest median pixel localisation errors for several trunk-associated landmarks (Supplementary Fig. 5c-e; neck, 3.58; left shoulder, 3.9; right shoulder, 4.03; back, 4.08). YOLO-CLS was trained using cropped images of each dog instance to achieve high individual-identification performance, reaching training and validation accuracies of 95% (Supplementary Fig. 4 and Supplementary Fig. 5f and g). In addition to trainings of YOLO models for dog detection and identification, YOLO-OBB model was trained to detect individual enrichment items. The model achieved a precision of 1.00 across all enrichment classes, while recall was 1.00 for all classes except toys (0.98; Supplementary Fig. 6).

**Fig. 3.**
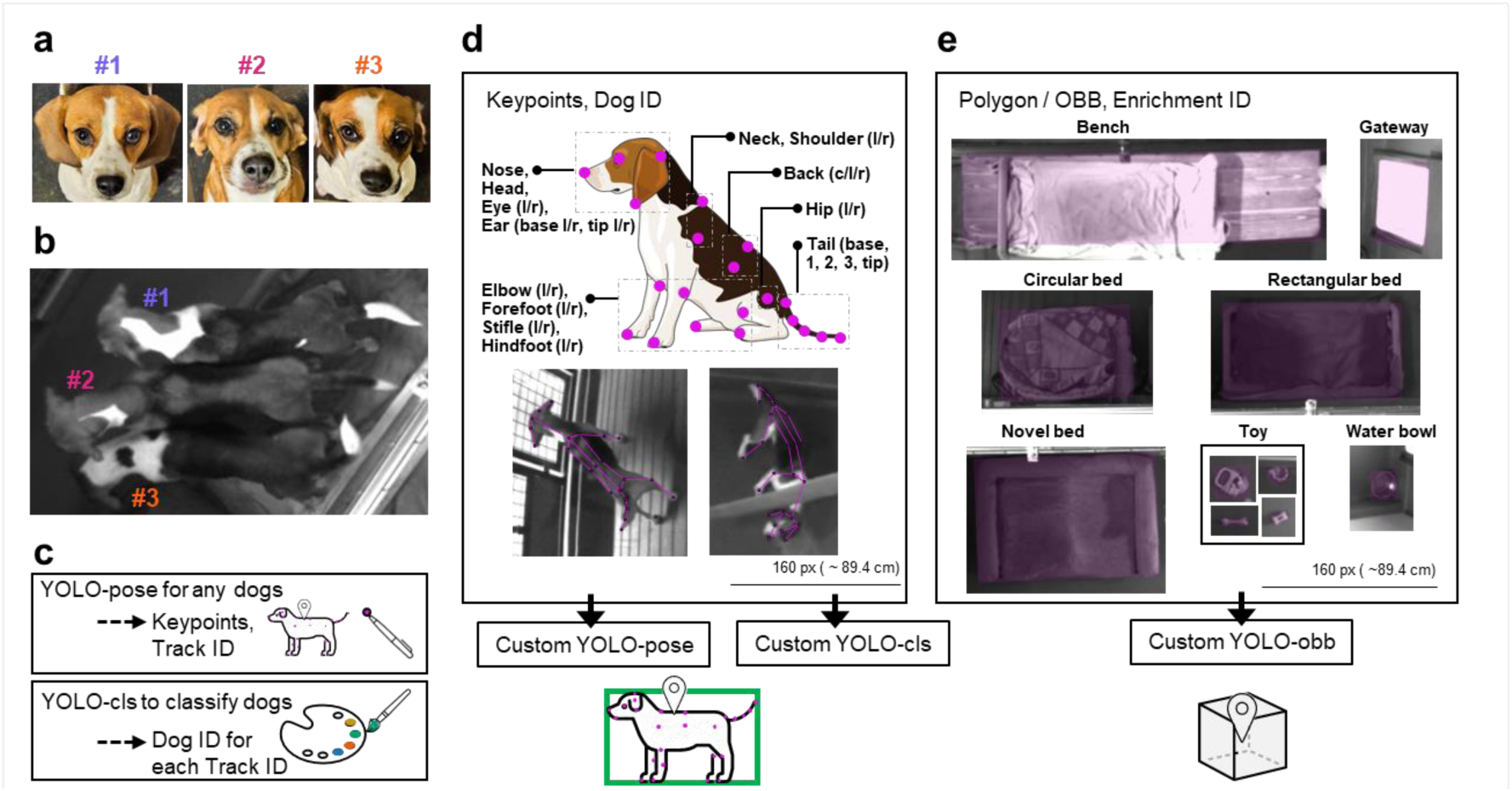
Analytical design with combination of CV models. **a)** Frontal view of female dogs. To visualise dog IDs, purple was assigned to dog 1, magenta to dog 2, and orange to dog 3. **b)** Dorsal view of female dogs. **c)** Key strategy to detect body parts with ID. Female d) 29 keypoints assigned for dog-pose model and a representative dog annotated with keypoints. e) Each enrichment item annotated with an oriented bounding box (OBB).

The generalisability of the identity-independent, female-trained YOLO-POSE model was evaluated on male dogs (Supplementary Fig. 5h). The model achieved an overall recall 85.2%, with 96.2% precision, mAP50 of 0.901, and an mAP50-95 of 0.554 (Supplementary Fig. 5i). Keypoint localisation errors were lowest for the right back (5.58), neck (6.27), left shoulder (7.64), right shoulder (8.52), and back (6.9), demonstrating transferability of the model to previously unseen individuals (Fig. 5j and k).

### Location tracking

Because kennel conditions followed highly reproducible daily routines (Fig. 1b), full-day recordings provided representative behavioural samples across recurring environmental and behavioural states.

Application of the trained YOLO models enabled tracking of each individual dog throughout the day (Fig. 4a-c). Although occupancy was generally spatially constrained, dogs exhibited broader area utilisation during active periods, most notably in the morning and afternoon, but also to a lesser extent during the night (Fig. 4b). Detection coverage exceeded 80% during periods when caretakers were absent and dogs were expected to remain inside the kennel (Fig. 4c), indicating robust and comparable tracking performance across individuals.

**Fig. 4.**
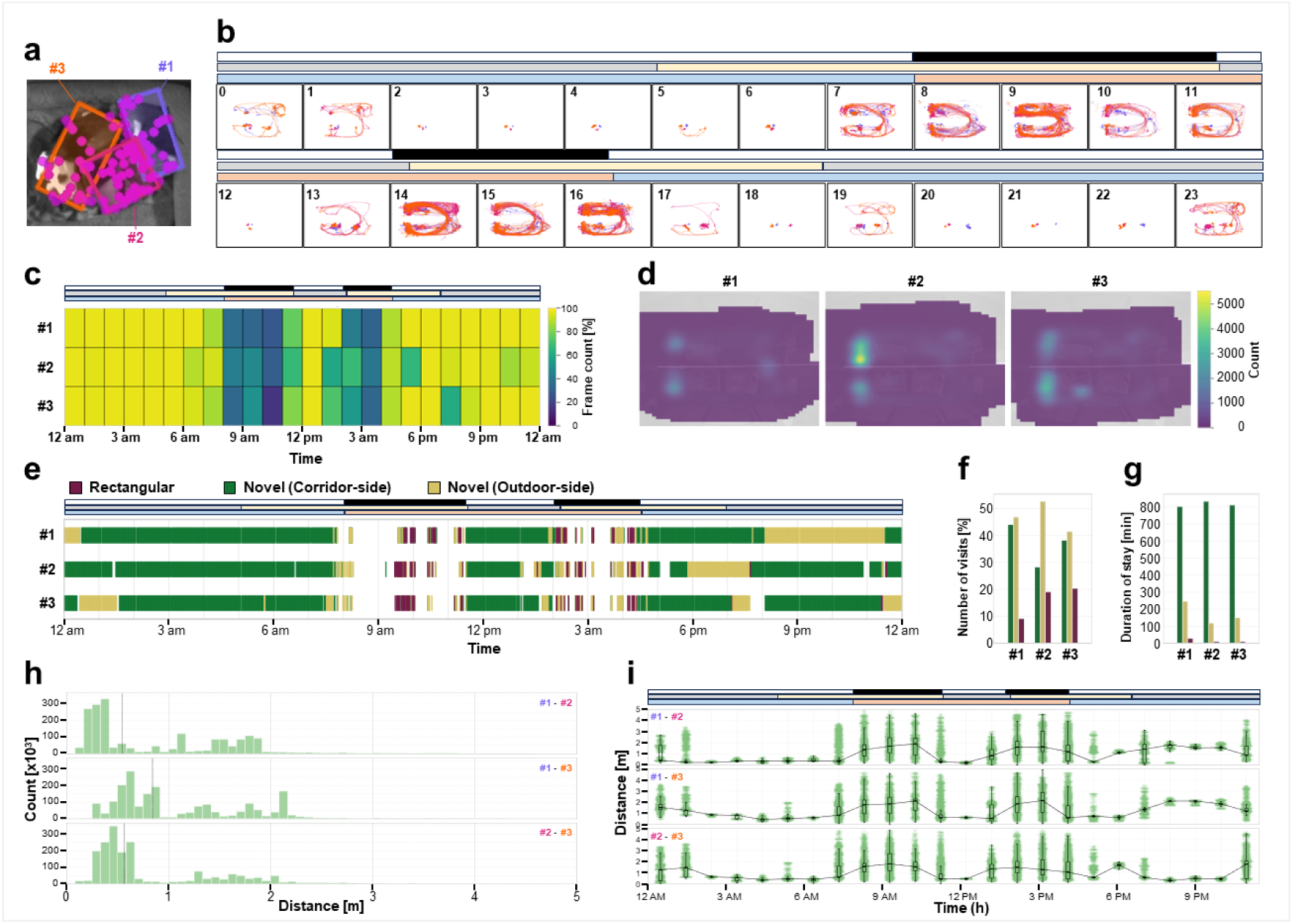
Hourly characteristics of dogs’ location on a representative day. **a)** Representative image of three female dogs detected in a bed with keypoints and OBB. **b)** Trajectory of each dog in each hour. **c)** Number of predictions per hour. **d)** Location heatmap outside the beds. **e-g)** Location characteristics inside the beds. Timeline of dogs’ present in beds (e). Number of times spent in each bed (f). Total time spent in each bed (g). **h-i)** Social distance between the dogs. Vertical lines indicate the median distance.

Aligning dog trajectories with enrichment detections enabled quantitative assessment of each dog dog’s location patterns outside and inside beds (Fig. 4 d and e). Heatmaps of the location of each individual outside the beds demonstrated distinct spatial occupancy patterns unique to each dog (Fig. 4d). Dog 1 occupied regions beneath the separation wall between kennel compartments, whereas Dog 2 predominantly remained near the upper corridor-facing upper corner of the kennel. Dog 3 exhibited broader spatial exploration with increased occupancy near the corridor in the lower region of the kennel. Analysis of inside-bed occupancy revealed that all dogs visited the newly introduced beds more frequently than the original rectangular bed type (Fig. 4e and f), while among the novel beds, the corridor-side bed exhibited the longest cumulative occupancy duration (Fig. 4g).

In addition to individual spatial occupancy, the dog trajectories enabled assessment of inter-individual relationships between all possible dog pairs (Fig. 4h and i). Dog 1 and Dog 3 consistently maintained the greatest separation (Fig. 4h), although pairwise distances fluctuated during active periods and became more stable during resting periods (Fig. 4i). Periods of reduced pairwise distances during night-time and daytime resting phases corresponded to the bed occupancy patterns shown in Fig. 4e (Fig. 4i).

Because kennel modifications for CV monitoring included the introduction of novel beds (Fig. 1a, Fig. 3e and Fig. 4e), bed occupancy was analysed over repeated recordings across 10 days to assess how environmental changes influenced resting-site selection. This analysis focused on bed type and location rather than individual differences between dogs. In the original kennel configuration, dogs frequently visited the circular bed located on the corridor side of the lower kennel compartment (Fig. 5c) but spent the longest cumulative time in the rectangular bed located on the outdoor side of the upper compartment (Fig. 5d). Following the installation of novel beds in the upper compartment, visitation frequencies became more evenly distributed across beds (Fig. 5e), whereas occupancy duration strongly favoured the newly introduced beds on both the corridor and outdoor sides of the upper compartment (Fig. 5f).

**Fig. 5.**
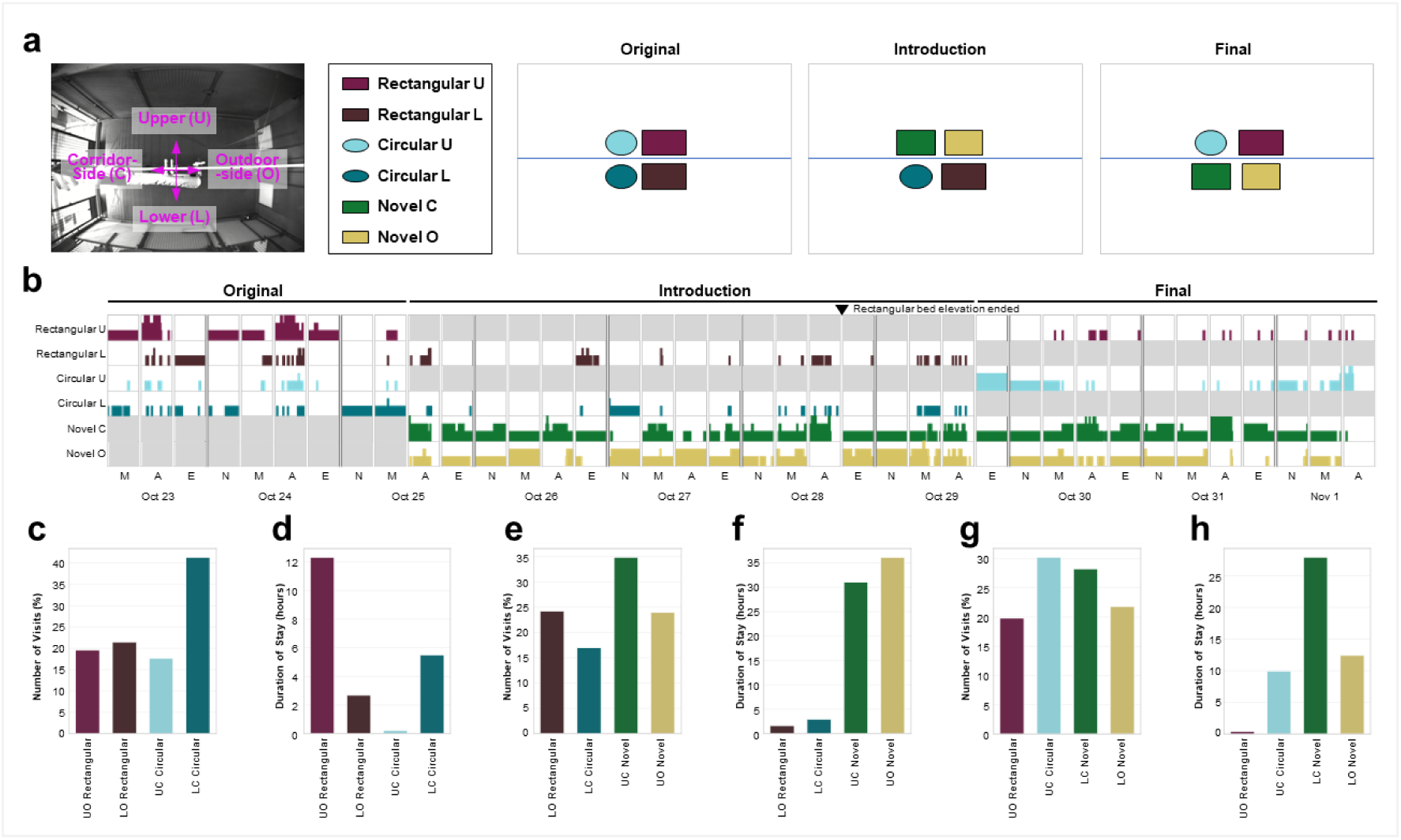
Bed occupancy across changes in bed type, position, and elevation. **a)** Schematic overview of bed arrangements. **b)** Temporal distribution of dogs interactions with each bed instance, shown as the number of dogs interacting with each bed in 1-min intervals across days and times of day. Grey boxes indicate periods when the corresponding bed instance was unavailable. M, morning; A, afternoon; E, evening; N. night. The triangle indicates the end of rectangular-bed elevation period. **c-h)** Quantification of bed occupancy across three experimental phases: original phases (c-d), introduction phase (e-f), and final phase (g-h). Bed occupancy was measured as the number of visits (c, e, g) and duration of stay (d, f, h), with values expressed as totals within each stage.

To further examine whether the preference toward upper compartment was associated with the preference toward the novel bed, bed locations were subsequently rearranged between the upper and lower compartments. Meanwhile, since the novel beds has elevation from the floor having legs, elevation of rectangular beds being placed on bench was finished at the end of the introductory phase. Notably, the novel bed located on the corridor side remained highly preferred of position. Interestingly, the circular bed exhibited higher occupancy than the rectangular bed (Fig. 5g, h). These changes in resting-site along the environmental rearrangement suggest that both bed design and kennel location contribute to preference in group-housed dogs.

### Locomotor activity

Beyond spatial distribution and inter-individual proximity, the tracked trajectories also enabled quantification of locomotor activity through frame-to-frame displacement of individual centroid positions. Activity patterns exhibited clear temporal variation, with strongly reduced movement during nocturnal resting periods (Fig. 6a, b). Dog 1 displayed consistently lower locomotor activity compared with Dogs 2 and 3. Over the 24-hour observation period, total travelled distance reflected these individual differences, with Dog 1 travelling 1.39 km, compared with 3.37 km for Dog 2 and 2.61 km for Dog3 (Fig. 6c).

**Fig. 6.**
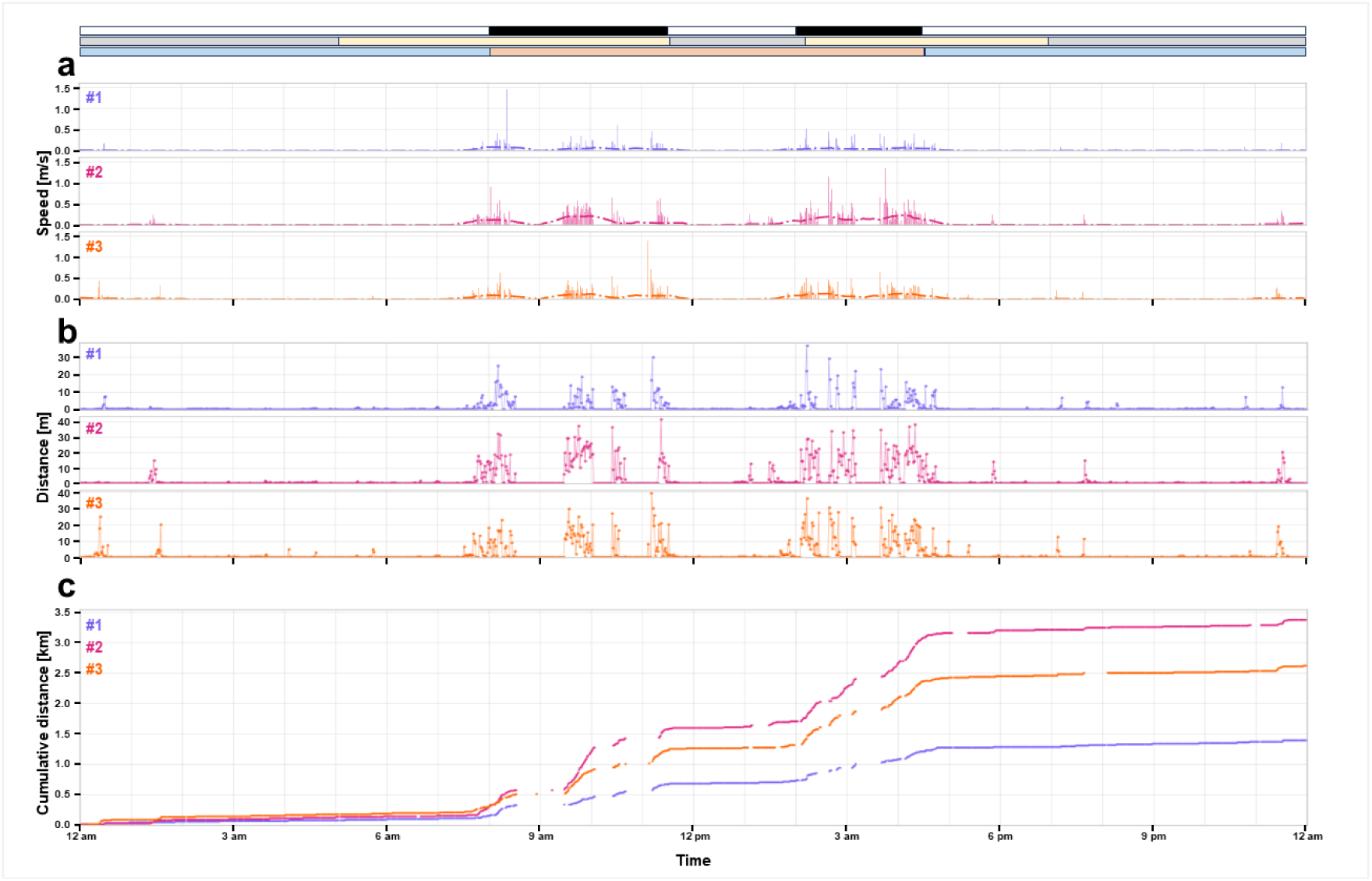
Locomotor activity profiles of individual dogs over a 24-hour period. **a)** Velocity. **b)** Locomotor activity. **c)** Cumulative travelled distance.

### Targeted interaction

While centroid trajectories enabled analysis of spatial occupancy, proximity, and locomotor patterns, keypoint detection provided finer-scale measurements of body orientation and specific interactions with environmental features.

Frequent occupancy near the corridor-facing side of the kennel suggested attention towards external activity and caretaker presence (Fig. 4d). Analysis of body orientation and distance from the corridor revealed that Dog 2 and Dog 3 are oriented towards the corridor more than twice as frequently as Dog1 (Fig. 7a-c), indicating marked inter-individual differences in potential human-directed behaviour.

**Fig. 7.**
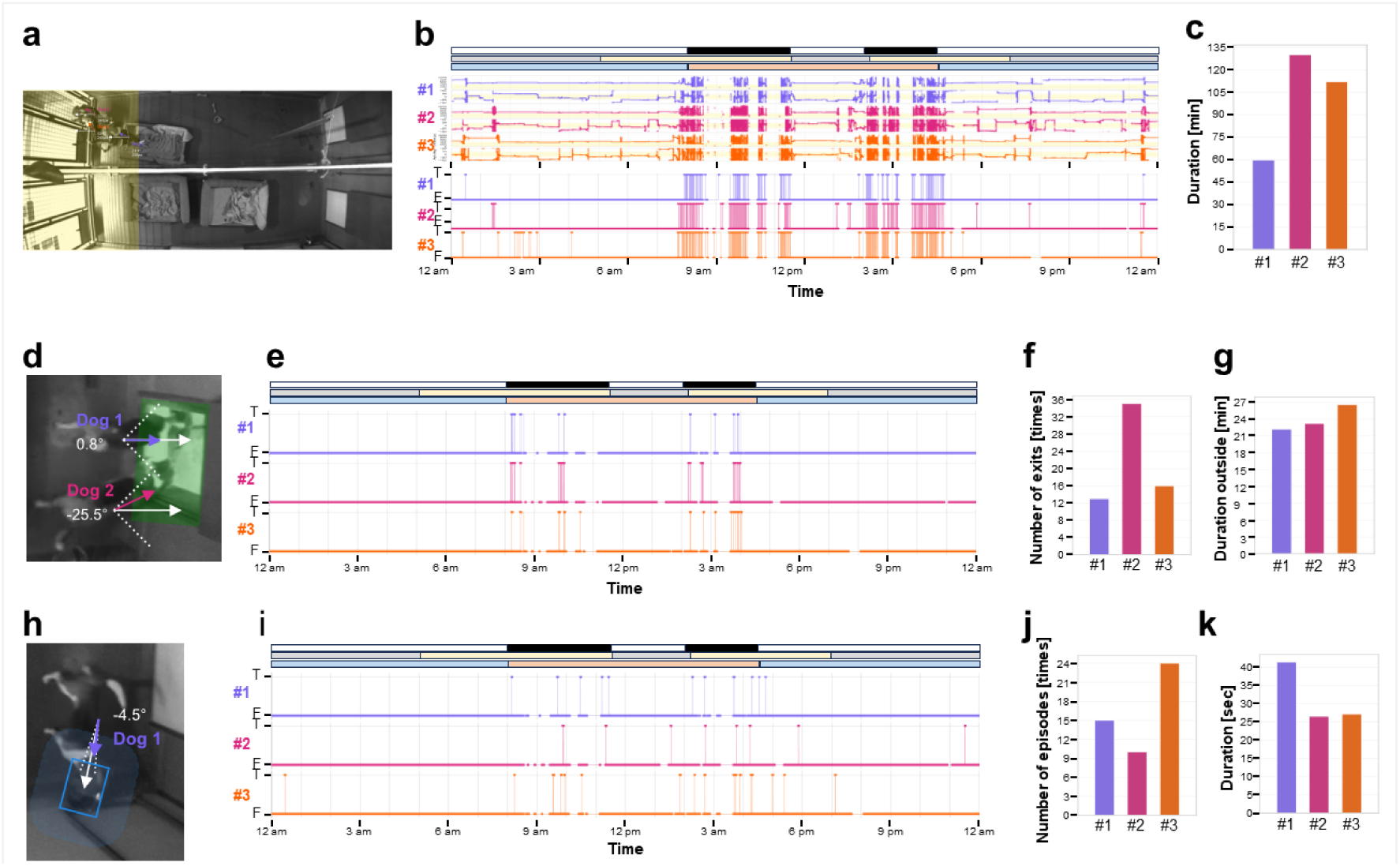
Behavioural analysis using keypoint predictions over a 24-hour observation period. **a-c)** Attention towards caretakers: representative frame of dogs at the corridor (a), timing of corridor visits (b), and total time spent at the corridor (c). **d-g)** Gateway use: representative frame of dogs using a gateway (d), timing of gateway use (e), number of exit episodes (d), and total duration spent outdoors (e). **h-k)** Water intake: representative frame of a dog drinking from a water bowl (h), timing of drinking events (i), number of drinking episodes (j), and total time duration of drinking (k).

Gateways to the outdoor kennel were opened twice daily by caretakers when they were present (Fig. 1b), allowing dogs to freely choose whether to access the outdoor area. Analysis of body orientation and location (Fig. 7d and e) revealed that Dog2 crossed the gateways more frequently than the other dogs (Fig. 7f), although total outdoor access duration remained comparable between individuals (Dog 1: 22 min; Dog 2: 23 min; Dog 3: 25 min; Fig. 7g).

Access to water bowls within the kennel was available throughout the day and night. Analysis of body orientation and distance from the water bowl enabled drinking events during both daytime and nocturnal periods (Fig. 7h). Dog 2 and Dog 3 accessed the water bowl during nighttime hours, with Dog 3 exhibiting the highest frequency (Fig. 7j). In contrast, Dog 1 spent the longest time near the water bowl while oriented toward it (41 min; Fig. 7k).

### Behavioural flow analysis

Sequential keypoint dynamics provide a framework for identifying differences in behavioural state without requiring behavioural labels. Behavioral flow analysis (BFA), originally developed for mice (Von Ziegler et al., 2024), was applied to different awakening periods (Fig. 8a), revealing distinct time-of-day-dependent behavioural patterns and differences in postural and locomotor dynamics between morning and afternoon awakening states (Fig. 8b-e).

**Fig. 8.**
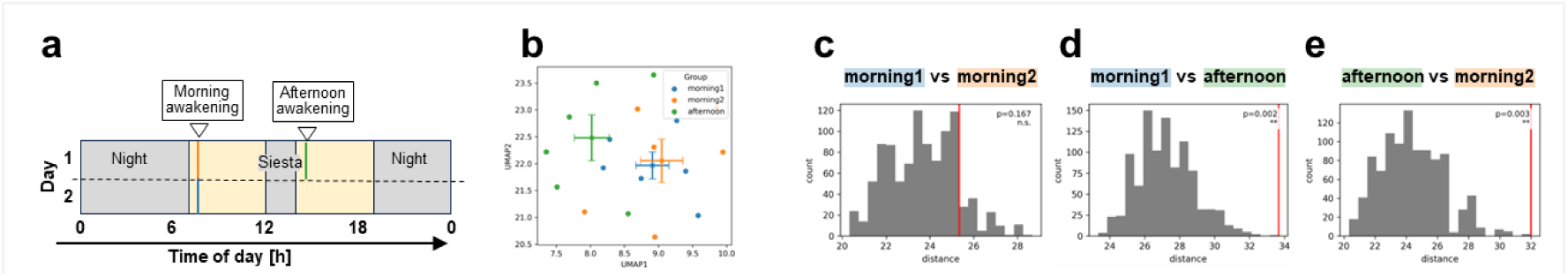
BFA. **a)** 24-hr schedule of dogs selected for BFA. Three 5-min awakening periods sampled either in the morning (Day 1: blue, Day 2: orange) or in the afternoon (Day 1: green). **b)** Behavioral flow fingerprinting (BFF). **c-e)** BFA reveals behavioural difference between two different awakening periods, but no significant differences between days.

## Discussion

In this study, we developed a modular CV framework for continuous behavioural profiling of group-housed dogs by integrating real-time YOLO architectures for pose estimation, Re-ID, and environmental object detection. Separating pose estimation from identity assignment enabled the YOLO-POSE model to generalise to previously unseen animals, allowing the framework to be adapted to new groups of animals by retraining only the YOLO-CLS model for Re-ID. The framework enabled quantification of spatial occupancy, locomotor activity, inter-individual proximity, and environmental interactions of each individual throughout the day and night under routine kennel conditions. Together, these findings demonstrate the feasibility of unbiased, scalable, non-invasive behavioural monitoring while extending CV beyond simple tracking towards quantitative behavioural phenotyping.

The YOLO-CLS model for Re-ID achieved 95% accuracy. Although Re-ID remains a challenging task in CV for behavioural analysis (Ravoor & T.S.B., 2020; Vidal et al., 2021), successful applications have been reported across several animal species, including tigers, giant pandas, chimpanzees, and brown bears (Liu et al., 2019; Schofield et al., 2019; Chen et al., 2020; Clapham et al., 2020). Dogs exhibit considerable variation in coat patterns (Brancalion et al., 2021), which may facilitate reliable visual identification. Compared with a previous study on group-housed Beagles that achieved similar performance only by colour-coding individuals with harnesses (Eberhardt et al., 2026), the present framework achieved comparable accuracy without physical markers. Nevertheless, validation across larger and more diverse populations will be necessary to determine the robustness and generalisability of the approach. Technical limitations remain, including vulnerability to occasional identity switching during close interactions, reduced performance under challenging lighting conditions, and the relatively small sample size.

Despite sharing the same housing environment, the dogs exhibited individual differences in spatial occupancy, activity, corridor-oriented behaviour, and resource use. Such variation highlights the potential value of individual-level assessment, as population-level summaries may obscure biologically relevant differences between individuals. Dogs are known to be highly sensitive to human attentional states and social cues (Call et al., 2003) and exhibit well-developed socio-cognitive abilities directed toward humans relative to other domestic species (Jardat & Lansade, 2022). The increased corridor-oriented behaviour observed in Dogs 2 and 3 may reflect greater interest in human activity and caregiver presence. However, alternative explanations, including general interest in corridor activity or anticipation of resource access, cannot be excluded.

Alignment of dog posture to environmental resources enabled quantification of evaluation of environmental use. Newly introduced beds accommodated dogs for more time over the original beds. Opportunities to express environmental preferences are recognised as an important component of environmental enrichment and welfare (Newberry, 1995; Wells, 2004) and previous studies have similarly reported that dogs exhibit preferences for particular resting substrates (Döring et al., 2018; Hubrecht et al., 1992). Rearrangement of bed locations demonstrated that environmental context influenced the selection of resting-site as well. Resource use therefore reflects the value animals assign to different components of their environment rather than simply their availability. This objective framework for evaluating environmental modifications and enrichment strategies from the animal’s perspective is consistent with calls for welfare assessment to incorporate animal-centred measures of environmental value and agency (Webber et al., 2022).

The distinct behavioural dynamics across awakening periods using BFA demonstrates the importance of continuous, data-driven behavioural phenotyping. Rather than relying on predefined observation periods or manually annotated behavioural categories, sequential keypoint dynamics can reveal temporal changes in behavioural state that may otherwise remain undetected.

Future development of this framework should focus on establishing validated, longitudinal, and individual-centred behavioural profiling approaches to advance objective assessment of animal behaviour and welfare. Longitudinal individual behavioural analysis will be required to establish baselines and detect deviations from individual-specific patterns over time. Such profiles may provide sensitive indicators of changes associated with husbandry and management interventions, health status, or environmental conditions. Larger cohort studies integrating behavioural data with physiological, health, and environmental measures will be important to validate the biological relevance of these behavioural signatures. While the present study focused primarily on behavioural profiling and resource use, future applications may include objective evaluation of enrichment strategies, husbandry interventions, training procedures, enclosure design, and veterinary management practices by quantifying how animals interact with and value different resources. Such approaches may support more evidence-based environmental design and refinement of animal care practices. However, the behavioural metrics identified in this study require further validation against detailed ethological observations and independent welfare indicators. Although behavioural data alone cannot directly measure emotional state, movement patterns, spatial behaviour, resource use, and temporal dynamics may provide valuable indicators for investigating relationships between behavioural variation and welfare-relevant processes. Integrating these behavioural features with automated classification approaches and emerging CV methods for recognising pain and postural cues (Veissier et al., 2019; Broomé et al., 2023) may further strengthen the welfare relevance of continuous monitoring frameworks. Collectively, continuous behavioural profiling has the potential to animal behaviour and welfare assessment from episodic observations towards longitudinal, individual-centred approaches, improving data quality, reducing reliance on manual observations, and supporting evidence-based refinement of animal care and management practices in line with the principle of the 3Rs.

## Conclusion

We present a modular CV framework that enables generalisable pose estimation, robust individual re-identification, and continuous behavioural profiling of group-housed dogs. Applied here to routine kennel conditions and a simple environmental modification, the framework demonstrates that individual differences in spatial behaviour, activity, resource use, and environmental engagement can be quantified objectively and continuously from standard video recordings. This work contributes to computational ethology and animal welfare science by providing a scalable, non-invasive approach for individual-centred behavioural analysis in complex group-housing environments. The framework is designed to support an expanding programme of research at the intersection of behaviour, welfare, and husbandry, with future applications including the longitudinal evaluation of housing interventions, environmental enrichment, and management practices. It has potential relevance not only for research settings but also for practical applications in veterinary monitoring, shelter management, and evidence-based refinement of animal care.

## Methods

### Animals and Housing

Seven Beagles (3 females littermates, 4 males of 2 pairs of 2 littermates) aged between 2-4 years were observed and housed at the faculty-owned research dog facility of the University of Zurich, Switzerland. Animals were group-housed (n = 2-4) in indoor kennels measuring either full (2.54 m x 4.47 m) or half size (1.27 m x 4.47 m). All dogs had *ad libitum* access to a paired outdoor kennel (2.74 m x 5.5 m) via rectangular gateways while caretakers were present. Caretakers entered the facility twice a day on weekdays to clean, feed, and train the dogs, and to allow access to the outdoor areas (Fig. 1b). Doors on the corridor were opened by caretakers occasionally to allow routine husbandry activities including temporarily removing individual dogs from their kennels for training sessions or to allow supervised play in the outdoor garden. In addition, dogs were walked 1-2 times per week or occasionally as part of research project. The facility was illuminated by natural sunlight through ceiling windows (24 windows; dimensions 1 m x 0.65 m) supplemented by artificial corridor lighting controlled by timers operating from 05:00 to 19:00 and adjusted for daylight saving time (Fig. 1b). Infrared projectors were automatically activated when ambient brightness dropped below a specific threshold. Water was provided *ad libitum* via water bowls and water nozzle. Feeding occurred once daily at 12:00, with additional snacks provided for training and enrichment. During the study, a strategic bedding transition was implemented; new beds were introduced to replace older benches. This change was necessary to follow animal housing regulations while eliminating visual occlusions caused by dogs hiding under elevated platforms.

The seven dogs included in this study were owned by the University of Zurich and were housed at the faculty-owned research dog facility under the animal husbandry license number #ZH191/0000 (Swiss authorization #255). All methods were carried out in accordance with relevant guidelines and regulations. No animals were euthanized, sacrificed, or killed as part of this study.

### Data Acquisition

Video recording was conducted using high-resolution cameras (Basler AG, Basler ace Classic acA1300-60gm) equipped with a lens (Computar, H0320KP) and a C-to-S mount adapter mounted to the ceiling at height of 3.10 m above the floor. The cameras were connected via switch (Ubiquiti, UniFi Switch Pro-48 PoE) to a computer equipped with a motherboard (MSI, Z790 GAMING PLUS WIFI), a processor (Intel, Core i9-14900KF, 24 cores, 32 logical threads, 3.2 GHz base frequency), a RAM 64 GB, a 2 TB internal solid-state drive (WD, Blue SN580), an 8 TB portable SSD external drive (Samsung, T5 EVO), an Ethernet network adapter (Intel, X710-DA4 converged). Recordings were triggered using either Pylon viewer manually or by Windows Task Schedular and some Python scripts. Videos were saved at 30 fps, 1280 × 960 pixels in either AVI or MP4 format. AVI videos were converted to mp4 format before analysis.

### Solar Events

Time of dawn, sunrise, sunset, and dusk at the facility location were obtained using the Astral Python package based on the facility’s geographic coordinates (Zurich, Switzerland; 47.497160° N, 8.652537° E) and local time zone (Europe/Zurich).

### MAE

To assess any change in frames, frames were paired sequentially, and MAE between each pair of frames were calculated from absolute difference of pixel intensity at each pixel. The resulting data enabled the identification of peak activity periods for each cage and recording session.

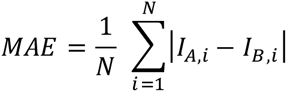

### Brightness Proxy

Average pixel intensity within a defined wall region on high wall position shown in Fig. 1c was calculated as brightness proxy.

### Door Status Analysis

To monitor the status of the kennel doors, a door-status analysis framework was implemented using a geometric approach (Supplementary Fig. 2). Prior to door-state detection, the lighting conditions in each frame were evaluated. Should the brightness fall below a predefined threshold, the lights were classified as being off. Given the assumption that door openings would not occur during periods of darkness (i.e. night-time and scheduled rest periods), door-state detection was suspended for that frame and all doors were assumed to a closed state. First, a single image frame that has optimal illumination and both doors fully closed was assigned as a primary reference frame for the kennel. Second, a ROI was defined for each door to include visual features that were informative for detecting changes in door position. Third, the reference image was cropped with a door-specific ROI, enhanced for the contrast using contrast-limited adaptive histogram equalization (CLAHE) (Zuiderveld, 1994), noise reduced applying a Gaussian blur with a 3 × 3 kernel, the edges were identified following Canny edge detection (Canny, 1986), and then the edges are dilated using a 3 × 3 kernel to enhance edge continuity. Finally, each image for the analysis was cropped and processed the same as the reference to obtain an enhanced binary edge mask, and assessed for the stability value by calculating the proportion of overlapping edge pixels between the current frame mask and the corresponding door-specific reference mask. High stability values indicated resemblance to the closed-door reference state, whereas lower values indicated a deviation consistent with door opening. A door was classified as true open when its stability value remained below the opening threshold for a specified number of consecutive candidate frames. Conversely, closure was detected when the stability value exceeded a dynamic closing threshold for a required number of frames. During each open episode, the closing threshold was dynamically adjusted relative to the mean stability observed whilst the door was open. To find a ROI that yields the best detection, several candidate ROIs covering each door were tested, and compared.

### Training YOLO models

Three YOLO models were selected to predict dogs and enrichments. For dogs, YOLO-POSE was chosen to predict body part, and then YOLO-CLS to identify each individual. For enrichments, YOLO-obb was chosen to predict and identify each enrichment. To train CV models, image frames were extracted from only female kennel across 12 days including both when the ceiling lights were on and off (Supplementary Fig. 3a). Initially, frames were collected from some specific time window with fixed time interval. Secondly, one day from each different furniture layout (original, introduction, and final) were chosen to collect unique frames relying on high MAE of 1 or 5 minutes of time interval on each day. All the frames were named systematically to ensure traceability to the original videos. All the annotations were made at annotation platform computer vision annotation tool (CVAT). Trainings were conducted in a Windows computer with Nividia CUDA RTX 4000.

For dogs, first, visible 29 body parts (nose, head, eyes, ears, neck, shoulders, back, hip, tail, elbows, forefeet, stifles and hindfeet) were annotated with each keypoint grouped with an ID. (Fig. 3d). Next, labels were exported in YOLO pose format. Second, cropped images were created from OBB around each group of instances with padding around. Third, to avoid distortion of training set, all the annotated image frames were sorted chronologically and paired sequentially regardless of different annotation sets, and pairs with low MAE were removed that caused low MAE (Supplementary Fig. 3b and Supplementary Fig. 4a, d and g). Note that the image removal was conducted in two ways: whole-size frame for pose model (Supplementary Fig. 3c), and each group of the trimmed images for classification model (Supplementary Fig. 4b, e and h). After filtering, a total of 742 annotated frames and 2,155 instances remained to train YOLO-POSE model (Supplementary Table 1). To prevent temporal leakage, the images were split in rotation chronologically dividing into training (70%), validation (15%), and test (15%) subsets. To train YOLO-POSE, all the class were merged into a single class (class 0) so that the trained model detected any dogs regardless of identity. The training dataset was augmented (imgsz=1280, kobj=1.5, pose=12, hsv_v=0.6, degrees=180, scale=0.9, perspective=0.002, fliplr=0.3, mosaic=0.5, erasing=0.3, rotate=180) and additional Albumentations transforms, including motion and Gaussian blur, Gaussian noise, CLAHE, and random brightness and contrast adjustments. The model was initialised from yolo11n-pose and trained for 600 epochs. To train YOLO-CLS, the training dataset was augmented (imgsz=416, kobj=1.0, hsv_h=0.0, hsv_s=0.0, hsv_v=0.2, degrees=45, translate=0.05, scale=0.9, perspective=0.001, fliplr=0.0, erasing=0.1) and additional Albumentations transforms, including motion and Gaussian blur, Gaussian noise, CLAHE, and rondom brightness and contrast adjustments. The model was initialised from yolo11n-cls and trained for 300 epochs.

For enrichments, they were labelled for each class initially with polygon and later with OBB (Fig. 3e). Labels were exported in either YOLO mask or YOLO OBB format depending on in which type the labels were made. In contrast to the dog datasets, no enrichment annotations were excluded due to the comparatively static nature of these objects. Annotation frequency per enrichment class ranged from 541 to 2,619 instances (Supplementary Table 1). The images were split in rotation chronologically dividing into training (70%), validation (15%), and test (15%) subsets. To train YOLO-obb, all polygon labels were converted to OBB and used together with OBB labels created at CVAT. Augmentation factors were adjusted (fliplr=0.5, rotate=180). Yolo11n-OBB was trained for 100 epochs.

### Inference pipeline

YOLO models were applied for every frame in sequence to keypoint coordinates and ID of each dog and OBB coordinate and class ID of each enrichment. For high-speed inference, best.pt of the trained YOLO-POSE was converted to TensorRT engines.

For dogs, first, the best.engine of the trained YOLO-POSE was applied to whole-frame size videos using model.track() using fine-tuned botsort tracker, and the predictions (video name, frame index, confidence of each instance, track id, x-y coordinates of each keypoint, confidence of each keypoint), additional information for each instance (centroid and OBB around each instance) was calculated, and the predicted information and the information withdrawn were saved as a data frame. Second, the track id was summarised, up to 10 frames of representative frames from each track using 5 different methods were created referring to the OBB with padding. Next, the representative images were classified applying the best.pt of trained YOLO-CLS, summarised and merged to the track summary data frame. Next, frames with conflict that are defined with more than one instance with common dog ID in a single frame was assessed, and all the instances that belong to shorter tracks in conflict were removed (Supplementary Fig. 8). Finally, the dog ID was merged to the keypoint data.

For enrichments, best.pt of trained YOLO-OBB was applied to the whole-frame size videos using model.track() using fine-tuned botsort tracker, and the predictions (video name, frame index, confidence of each instance, track id, the enrichment name of each instance).

### Testing male keypoints

To assess the generalisability of the keypoint detection model, 32 independent image frames containing four male dogs in a male kennel were manually annotated using the same keypoint configuration as for the female dogs. These images were analysed using the inference pipeline developed for the female dogs, including the female-trained YOLO-POSE model. Individual identification with the trained YOLO-CLS was not performed. Predicted keypoints were matched to manually annotated ground-truth instances using intersection over union, and keypoint prediction accuracy was evaluated for the matched detections.

### Behavioural feature extraction

To extract behavioural features from the predicted dog poses, spatial, temporal, and interaction-based metrics were computed from the frame-wise tracking data, using only detections above the confidence threshold of 0.5. Furthermore, enrichment detections were incorporated to determine enrichment-related interactions and behavioural tendencies. A dog was classified as interacting with an enrichment if the defining body point was located inside the OBB of the enrichment. For most enrichments, the neck keypoint was used to represent directed attention or resource use. For beds, however, the midpoint between the neck and back keypoints was used to account for dogs resting inside the bed while extending their head outside the bed area. For analyses requiring body orientation, the neck and back keypoints were selected based on the high prediction accuracy and reliability they exhibited, and the fact that they formed a stable body axis for estimating the direction of the dog. The extracted behavioural features included bed preference, locomotor activity, social distance, location occupancy, detection frequency, corridor attention, gateway usage, and water intake across a 24-hour recording period.

To analyse bed occupancy, bed tracks were categorised into three bed instances for the 24-hour analysis based on bed type and spatial location: one old rectangular bed and two newly introduced beds. The two new beds were separated into left and right instances using the average OBB centroid position across each track duration, followed by splitting tracks at the median x-position. The multi-day analysis distinguished six bed instances, comprising upper and lower old rectangular beds, upper and lower old circular beds, and the left and right new beds. Interaction events were grouped into 1-minute intervals. In the 24-hour analysis, bins were computed separately for each dog and bed instance, whereas the multi-day analysis was independent of dog identity and counted a bed interaction if any dog interacted with the bed during the respective interval. Consecutive interaction frames were grouped into single visits, allowing gaps of up to 3 seconds. In contrast to the 30-fps employed for the 24-hour recordings, the multi-day recordings were processed at 1 fps.

To quantify social distance between dogs, the frame-wise Euclidean distance between tracked centroids of for all dog pairs was calculated. Pixel values were converted to metres using a conversion factor derived from the known kennel length and the corresponding pixel distance along the central kennel separation line. The distances were then grouped into hourly bins and visualised using scatter plots, boxplots, and median lines. Furthermore, the overall distance distributions across the 24-hour period were summarised as histograms, with the median distance indicated.

To quantify locomotor activity, centroid displacement was measured over 60-frame intervals, corresponding to approximately 2 seconds at 30 fps. Pixel displacement was converted to metres and divided by elapsed time to obtain speed in metres per second. Speed estimates were averaged within 1-minute bins and visualised as bar plots together with a smoothed mean curve using a 10-minute rolling window.

To quantify attention towards the corridor, a frame-wise binary classification was applied to determine whether dogs were attending to the corridor. A dog was classified as attending to the corridor when the neck and back keypoints were located within the corridor-associated area and the body axis, defined as the vector from the back to the neck keypoint, was oriented towards the corridor. Orientation towards the corridor was quantified as the angular deviation between the body axis and the inverted x-axis, representing the corridor direction within the kennel, with a threshold of ±75°. The aforementioned criteria were furthermore visualised as continuous measures, incorporating the distance from the left border of the corridor area and the angular deviation of the dogs from the direction of the corridor. The yellow regions in the figure delineate the threshold ranges that fulfil the attention criteria, and data points falling within these regions were classified as corridor attention.

To detect water intake from the water bowls a comparable frame-wise classification method was used. Dogs were classified as drinking if the neck keypoint was situated within the water bowl OBB, or within a 30-pixel buffer surrounding it, and if the body axis was oriented towards the bowl centroid within ±8°. Consecutive positive frames were grouped into drinking segments, with short interruptions of up to one second being permitted. The total duration of drinking was derived from the sum of all positively classified frames.

To identify usage of the gateway, the same frame-based classification approach was employed. Frames were classified as positive if either the neck or back keypoint was situated within the identified gateway OBB, and if the body axis was oriented towards the gateway within a range of ±45°. To eliminate false positives from dogs approaching a closed gateway, only segments followed by a minimum of 55 consecutive frames without detections were retained, as this indicated that the dogs had indeed gone outside. An additional buffer of 19 frames was permitted following each retained event to account for brief residual detections. Consecutive gateway frames were grouped into single exit events, while allowing short interruptions of up to one second.

To generate detection heatmaps that illustrate the frequency of reliable centroid detections across the 24-hour observation period, detections were grouped into hourly bins and expressed relative to the maximum possible number of frames.

To calculate cumulative distance, centroid positions were sampled at one-second intervals in the context of locomotor analysis. Euclidian displacement between consecutive positions was converted to metres, summed over time, and reduced to 1-minute resolution.

To generate activity heatmaps, which assesses spatial occupancy outside of the beds, frames that contained bed interactions were excluded, and centroid positions were accumulated across the entire recording period. These cumulative heatmaps were then subjected to a smoothing process using a Gaussian filter, with the aim of visualising the regions of high and low activity.

### BFA

x and y coordinates of three body parts (head, back and tail set) of all dogs predicted with trained CV models were used to generate a set of features. The features were collected from 15 pre- and post-frames for each frame. For the detail, described in our previous paper (Von Ziegler et al., 2024). These patterns were used for BFF.

## Data availability

The data supporting the findings of this study are available upon reasonable request from the corresponding author.

## Funding Declaration

This research was supported by Refinement Grant 2023 (RG-2023-011) from Swiss 3R Competence Center, GRC Career Grant (2025_Q3_CG_004) from Graduate Campus at the University of Zurich, and financial support for 3R and Animal Welfare Projects from the Office for Animal Welfare and 3R at the University of Zurich.

## Abbreviations

BBOX: bounding box
BFA: behavioral flow analysis
BFF: behavioral flow fingerprinting
C: closed
CLAHE: contrast-limited adaptive histogram equalization
CLS: classification
CV: computer vision
CVAT: computer vision annotation tool
DLC: DeepLabCut
fps: frame per second
ID: identity
MAE: mean absolute error
mAP: mean average precision
O: open
OBB: oriented bounding box
Re-ID: re-identification
ROI: region of interest
SLEAP: social LEAP estimates animal poses
YOLO: You Only Look Once

## Acknowledgements

We thank the animal care staff of the research dog facility for their continuous support, dedicated animal husbandry, and assistance with daily kennel management throughout the study. Their commitment to high animal welfare standards and consistent care was essential for the successful implementation of the longitudinal behavioural monitoring system.

## Author contributions

O.S. and H.R. conceived an initial idea of the project.

H.R., T.L., M.S., O.S., L.I., D.N., and P.V.P. installed infrastructure for video recordings.

T.L. and D.N. wrote scripts or application to conduct video recording systematically.

H.R., M.S., and A.C. planned recordings.

M.S. and K.A. performed MAE analysis.

M.S. and P.V.P. extracted frames for data annotation.

M.S., P.V.P., J.G., K.A., P.U. and L.I. annotated image frames.

M.S., P.U., P.V.P. and J.G. designed analytical pipeline and trained CV models.

M.S. ran predictions.

T.L. wrote the scripts for BFA.

M.S., P.U., J.G., K.A. and P.V.P. analysed the data.

M.S., A.C., P.U., K.A. and H.R. wrote the manuscript.

H.R., O.S., and M.S. secured the fundings.

H.R. supervised the project.

All authors reviewed and agreed on the final manuscript.

## Competing interests

We have no conflicts of interest to disclose. All authors declare that they have no conflicts of interest.

## Notes

### Competing Interest Statement

The authors have declared no competing interest.

