## Supplemental Information for "Longitudinal computer vision-based behavioural phenotyping of group-housed dogs for translational and welfare research"

### Supplementary Information

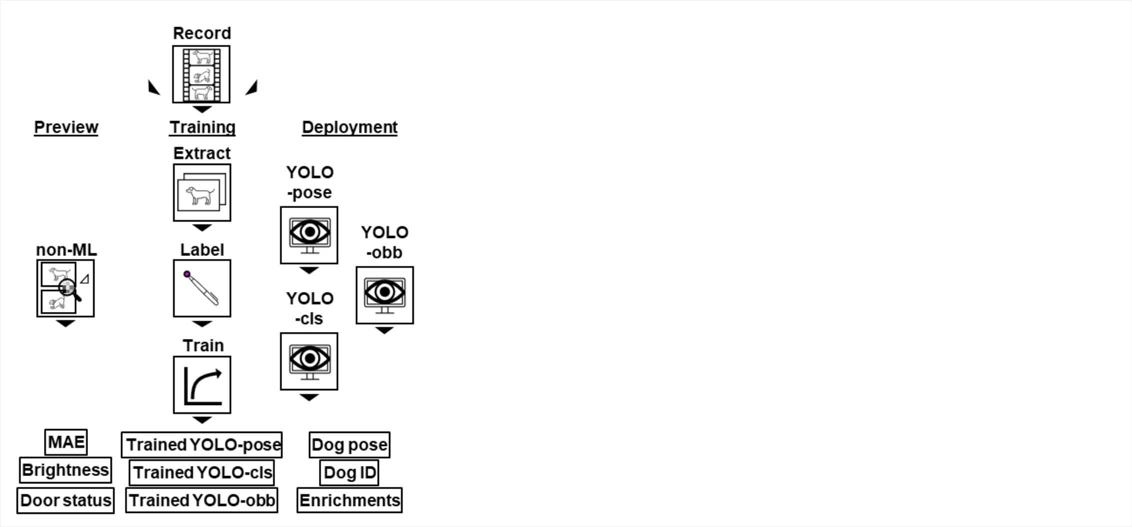


**Supplementary Fig. 1 | Workflow of video analysis.** The workflow consisted of three stages: preview, training, and deployment. The preview stage included non-machine learning (ML) computer vision (CV) analyses, including mean absolute error (MAE), brightness, and door status. During training, extracted frames were labelled and used to train YOLO-POSE, YOLO-CLS, and YOLO-OBB models. During deployment, the trained models predicted dog keypoints, individual identity, enrichment identity (ID), and location.

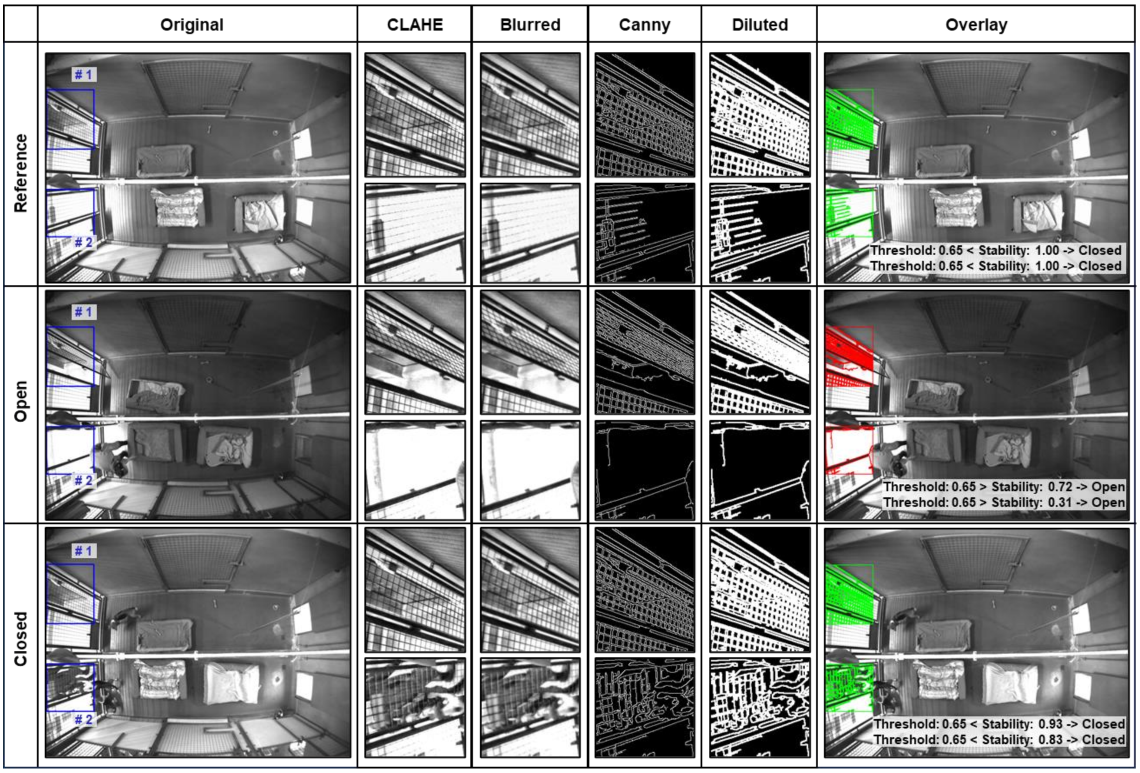


**Supplementary Fig. 2 | Automated detection of kennel door status.**
Blue boxes indicate the regions of interest (ROIs) used to analyse kennel door status. Cropped door ROIs were processed using contrast-limited adaptive histogram equalization (CLAHE), Gaussian blur, Canny edge detection, and dilation. Door status

was classified by comparing overlap between edge masks from reference and analysed frames. Green and red overlays indicate closed- and open- door status, respectively.

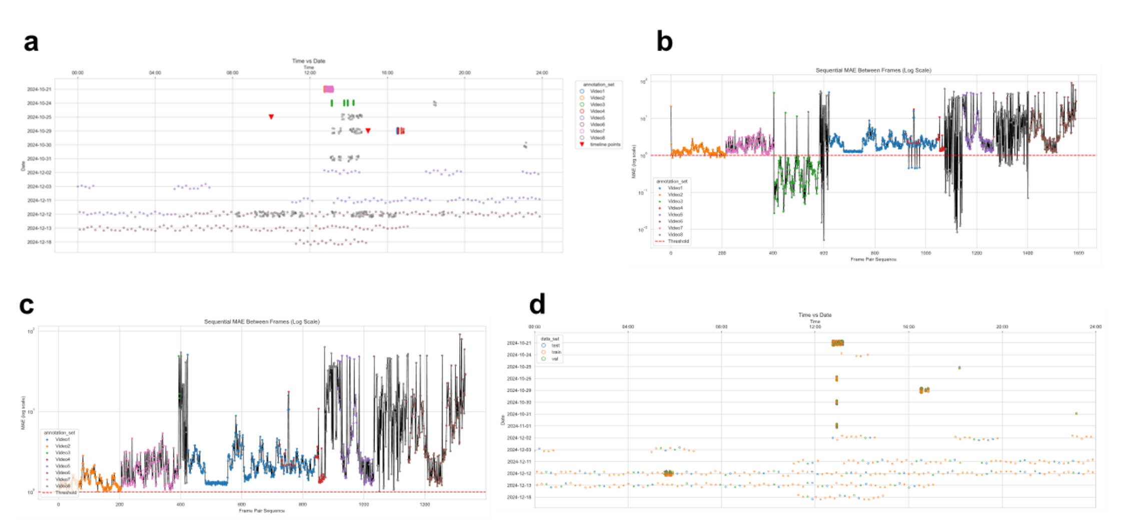


**Supplementary Fig. 3 | Temporal distribution and whole-frame MAE of annotated frames.**

**a)** Temporal distribution of all annotated frames. **b)** Whole-frame MAE between consecutive annotated frames. **c)** Whole-frame MAE of frames retained after high-MAE filtering. **d)** Temporal distribution of retained after high-MAE filtering. For a-c, points are coloured by annotation task set. For d, points are coloured by training date set.

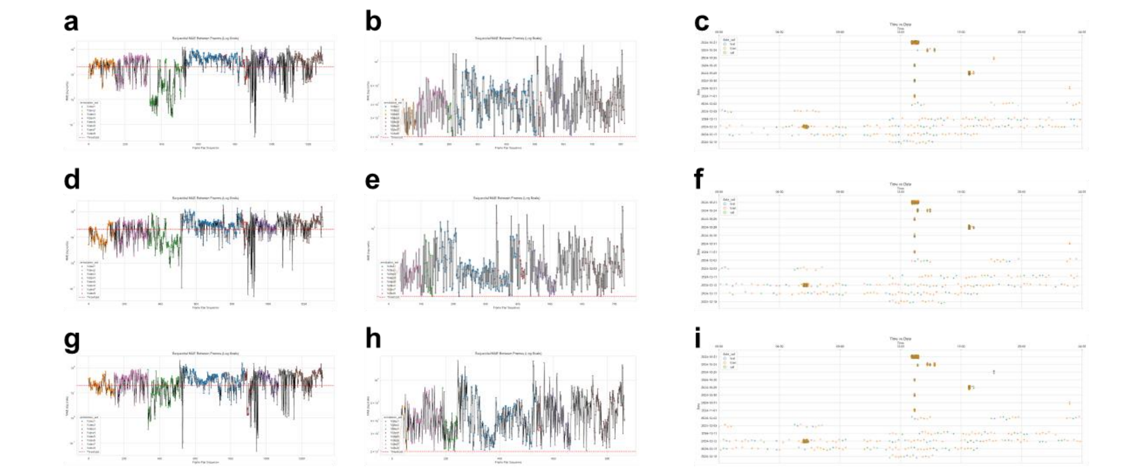


**Supplementary Fig. 4 | Dog-specific temporal distribution and cropped-frame MAE for YOLO-CLS.**

**a)** Cropped-frame MAE between consecutive images for Dog 1. **b)** Cropped-frame MAE of images retained after high-MAE filtering for Dog 1. **c)** Temporal distribution of retained images for Dog 1. **d–f)** Corresponding analyses for Dog 2. **g–i)** Corresponding analyses for Dog 3. For a, b, d, e, g and h, points are coloured by annotation task set. For c, f and i, points are coloured by training date set.

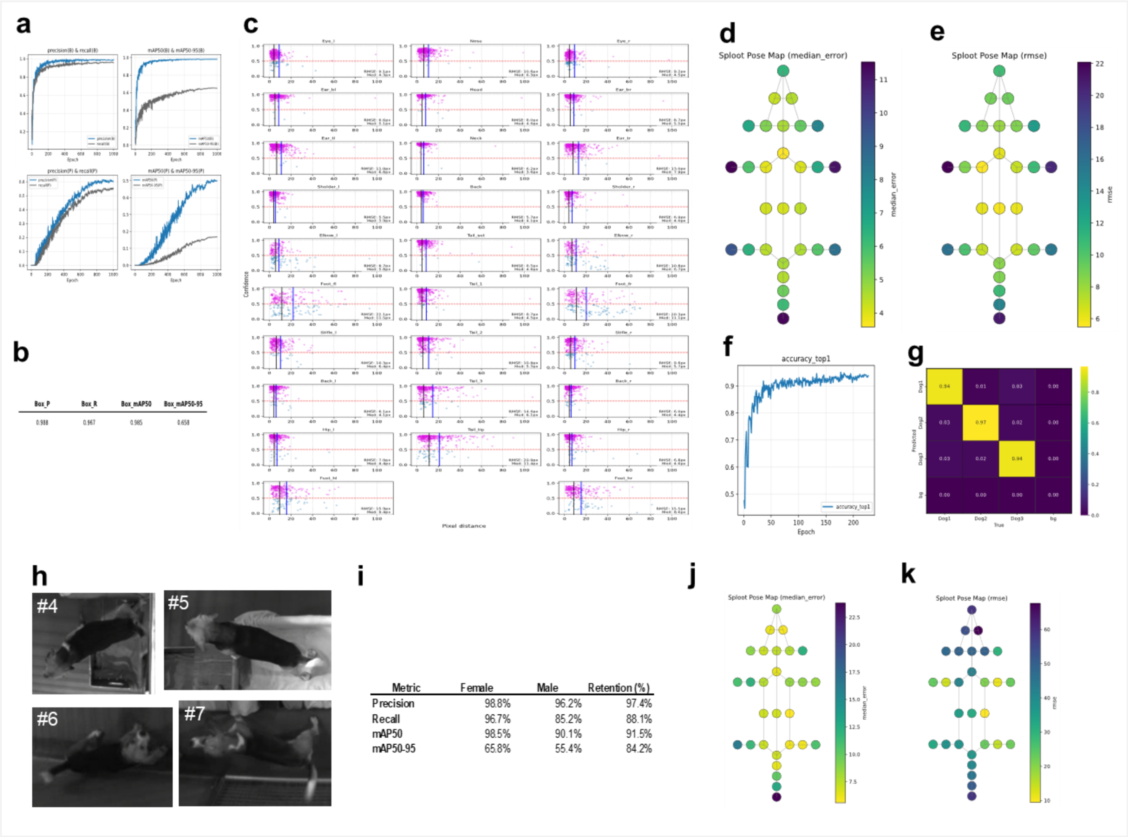


**Supplementary Fig. 5 | Performance of trained models for dog detection and identification.**

**a-e)** Performance of the trained YOLO-POSE model for keypoint estimation. (a) Training curves showing model performance across epochs. (b) Evaluation metrics on the test dataset. (c) Distribution of confidence versus error between predicted and ground truth. (d-e) Heatmaps of keypoint localisation error across body parts, with colours representing summarised pixel error between predicted and ground truth keypoints (d, median error; e, root mean square error (RMSE)).

**f and g)** Performance of the trained YOLO-CLS model for individual dog identification. (f) Training and validation learning curves across epochs. (g) Confusion matrix showing classification performance across individual dogs and the background class.

**h-k)** Performance of the trained YOLO-POSE model on unseen dogs. (h) Example dorsal images of male dogs with predicted keypoints. (i) Comparison of pose estimation metrics between male and female dogs. (j-k) Heatmaps of keypoint localisation error across body parts, with colours representing summarised pixel error between predicted and ground truth keypoints (j, median error; k, RMSE).

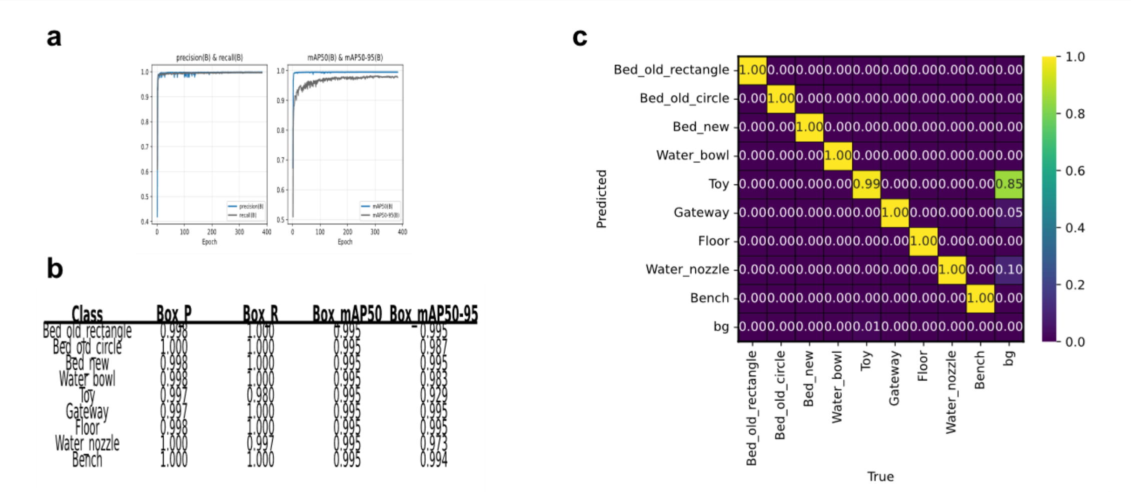


| Class | Box P | Box R | Box mAP50 | Box mAP50-95 |
|---|---|---|---|---|
| Bed old rectangle | 0.998 | 1.000 | 0.995 | 0.995 |
| Bed old circle | 1.000 | 1.000 | 0.995 | 0.987 |
| Bed new | 0.998 | 1.000 | 0.995 | 0.995 |
| Water bowl | 0.998 | 1.000 | 0.995 | 0.983 |
| Toy | 0.997 | 0.980 | 0.995 | 0.929 |
| Gateway | 0.997 | 1.000 | 0.995 | 0.995 |
| Floor | 0.998 | 1.000 | 0.995 | 0.995 |
| Water nozzle | 1.000 | 0.997 | 0.995 | 0.973 |
| Bench | 1.000 | 1.000 | 0.995 | 0.994 |

**Supplementary Fig. 6 | Accuracy of predictions of enrichment object segmentation.** **a)** Training and evaluation curves (loss, accuracy, mAP) for enrichment object segmentation. **b)** Precision–recall curves illustrating per-class performance for enrichment objects. **c)** Confusion matrix across enrichment object classes.

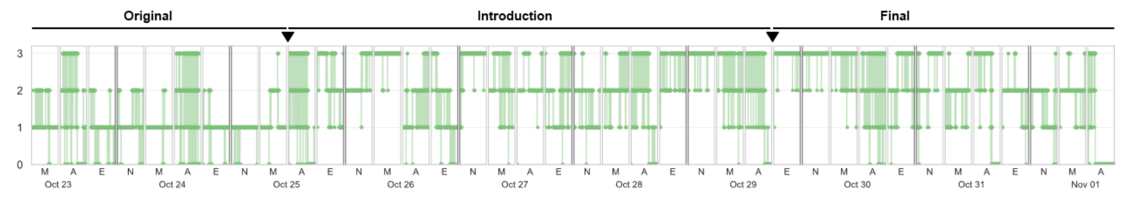


**Supplementary Fig. 7 | Number of dogs detected.**

Number of dogs detected at each time point, ranging from zero to three. The x-axis shows the recording timeline from 23 October to 1 November, divided into morning (M), afternoon (A), evening (E), and night (N) periods for each day. The y-axis indicates the number of detected dogs.

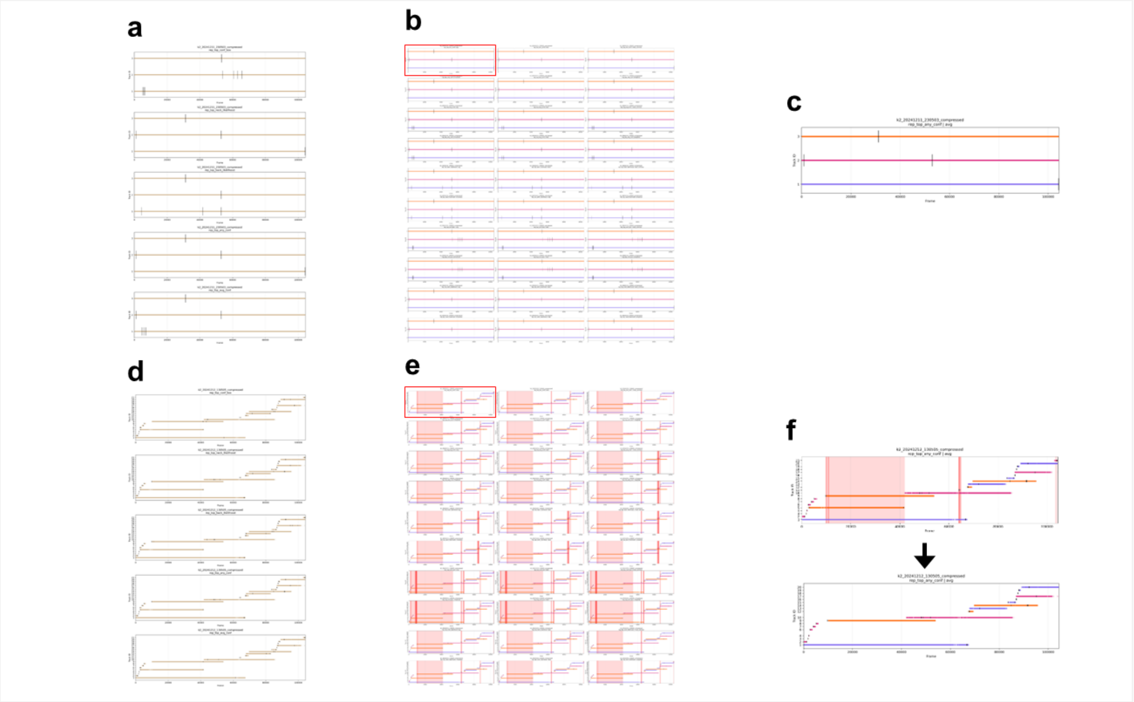

**Supplementary Fig. 8 | Track and ID summary.**
Representative summary of no ID conflict (a-c) and ID conflict in some frames (d-f). **a-c)** show no more than three tracks, corresponding to three individual dog IDs, which remained consistent throughout. **d-e)** show examples with more than three tracks per analysed sequence, requiring correction due to fragmented tracks and ID conflicts. Red shaded areas indicate frames affected by overlapping track ID conflicts. **f)** shows before and after resolving conflict of two parallel dog 3 tracks occurring at the same time.

| model | class | images | | | instances | | |
|---|---|---|---|---|---|---|---|
| | | train | val | test | train | val | test |
| YOLO-pose | Dog | 742 | 185 | 186 | 2’155 | 539 | 543 |

| model | point | images | | | instances | | |
|---|---|---|---|---|---|---|---|
| | | train | val | test | train | val | test |
| YOLO-pose | nose | 719 | 181 | 182 | 1’643 | 421 | 425 |
| YOLO-pose | head | 740 | 185 | 183 | 1’999 | 507 | 503 |
| YOLO-pose | neck | 742 | 184 | 184 | 2’011 | 501 | 503 |
| YOLO-pose | back | 740 | 184 | 184 | 1’934 | 479 | 487 |
| YOLO-pose | tail_set | 741 | 185 | 183 | 1’880 | 468 | 481 |
| YOLO-pose | tail_1 | 722 | 178 | 181 | 1’760 | 438 | 448 |
| YOLO-pose | tail_2 | 722 | 179 | 182 | 1’766 | 442 | 450 |
| YOLO-pose | tail_3 | 708 | 179 | 178 | 1’684 | 420 | 430 |
| YOLO-pose | tail_tip | 669 | 173 | 171 | 1’561 | 395 | 401 |
| YOLO-pose | Shoulder_l | 734 | 183 | 184 | 1’798 | 445 | 451 |
| YOLO-pose | Elbow_l | 509 | 128 | 131 | 744 | 184 | 197 |
| YOLO-pose | Foot_fl | 558 | 143 | 148 | 882 | 221 | 240 |
| YOLO-pose | Hip_l | 720 | 178 | 184 | 1’651 | 413 | 423 |
| YOLO-pose | Thigh_l | 659 | 162 | 167 | 1’135 | 300 | 285 |
| YOLO-pose | Foot_hl | 569 | 142 | 150 | 1’133 | 271 | 293 |
| YOLO-pose | Shoulder_r | 736 | 183 | 180 | 1’883 | 467 | 459 |
| YOLO-pose | Elbow_r | 546 | 129 | 126 | 787 | 189 | 188 |
| YOLO-pose | Foot_fr | 600 | 151 | 154 | 981 | 237 | 246 |
| YOLO-pose | Hip_r | 718 | 177 | 180 | 1’693 | 413 | 436 |
| YOLO-pose | Thigh_r | 660 | 162 | 167 | 1’237 | 296 | 311 |
| YOLO-pose | Foot_hr | 596 | 143 | 151 | 1’194 | 273 | 294 |
| YOLO-pose | Ear_bl | 739 | 185 | 183 | 1’933 | 492 | 483 |
| YOLO-pose | Ear_tl | 729 | 185 | 181 | 1’750 | 460 | 452 |
| YOLO-pose | Ear_br | 732 | 185 | 183 | 1’922 | 480 | 477 |
| YOLO-pose | Ear_tr | 728 | 183 | 181 | 1’758 | 433 | 431 |
| YOLO-pose | Eye_l | 680 | 178 | 175 | 1’408 | 367 | 362 |
| YOLO-pose | Eye_r | 667 | 168 | 168 | 1’239 | 320 | 314 |
| YOLO-pose | Back_l | 731 | 180 | 185 | 1’757 | 434 | 441 |
| YOLO-pose | Back_r | 730 | 182 | 179 | 1’788 | 443 | 455 |

| model | class | images | | |
|---|---|---|---|---|
| | | train | val | test |
| YOLO-cls | Dog1 | 543 | 136 | 136 |
| YOLO-cls | Dog2 | 488 | 122 | 122 |
| YOLO-cls | Dog3 | 577 | 144 | 144 |

| model | class | images | | | instances | | |
|---|---|---|---|---|---|---|---|
| | | train | val | test | train | val | test |
| YOLO-obb | Bed_new | 518 | 128 | 127 | 1’035 | 256 | 254 |
| YOLO-obb | Bed_old_circle | 576 | 144 | 144 | 841 | 210 | 210 |
| YOLO-obb | Bed_old_rectangle | 783 | 195 | 193 | 1’048 | 261 | 259 |
| YOLO-obb | Bench | 276 | 69 | 69 | 541 | 135 | 135 |
| YOLO-obb | Floor | 788 | 195 | 194 | 788 | 195 | 194 |
| YOLO-obb | Gateway | 788 | 195 | 194 | 1’576 | 390 | 388 |
| YOLO-obb | Toy | 784 | 195 | 194 | 2’619 | 651 | 641 |
| YOLO-obb | Water_bowl | 521 | 128 | 127 | 821 | 203 | 202 |
| YOLO-obb | Water_nozzle | 653 | 162 | 161 | 1’306 | 324 | 322 |

**Supp. Table1 | Labels used for training and validation.**

Summary of annotated image and instance numbers used to train, validate, and test the YOLO-POSE, YOLO-CLS, and YOLO-OBB models. The table shows the total dog class annotations for YOLO-POSE, as well as the distribution of individual keypoint labels across the training, validation, and test datasets. For YOLO-CLS, the number of images for each dog identity class are shown. For YOLO-OBB, image and instance numbers are listed for each annotated enrichment class.
